# Deciphering the RNA recognition by Musashi-1 to design protein and RNA mutants for in vitro and in vivo applications

**DOI:** 10.1101/2024.10.24.619864

**Authors:** Anna Pérez-Ràfols, Guillermo Pérez-Ropero, Linda Cerofolini, Luca Sperotto, Joel Roca-Martínez, Rosa Anahí Higuera-Rodríguez, Pasquale Russomanno, Wolfgang Kaiser, Wim Vranken, Helena Danielson, Alessandro Provenzani, Tommaso Martelli, Michael Sattler, Jos Buijs, Marco Fragai

## Abstract

RNA Recognition Motifs (RRMs) are essential post-transcriptional regulators of gene expression in eukaryotic cells. The Human Musashi-1 (MSI-1) is an RNA-binding protein that recognizes (G/A)U_1-3_AGU and UAG sequences in diverse RNAs through two RRMs and regulates the fate of target RNA.

Here, we combined structural biology and computational approaches to analyse the binding of the RRM domains of human MSI-1 with single-stranded and structured RNAs ligands. We used our recently developed computational tool RRMScorer to design a set of mutants of the MSI-1 protein to bind novel RNA sequences to alter the binding selectivity. The *in-silico* predictions of the designed protein-RNA interactions are assessed by NMR and SPR. These experiments also are used to study the competition of the two RRM domains of MSI-1 for the same binding site within linear and harpin RNA. Our experimental results confirm the *in-silico* designed interactions, thus opening the way for the development of new biomolecules for in vitro and in vivo studies and downstream applications.

## Introduction

The fate and regulation of mRNAs are controlled by RNA-Binding Proteins (RBP), which recognize specific oligonucleotide sequences non-covalently. RBPs can have single or multiple domains responsible for the recognition and binding of RNA. The RNA Recognition Motif (RRM) ^1,2^ is a well-studied and widespread RNA-binding domain in RBP in higher vertebrates ^3^. The RRM domain comprises about 90 residues and adopts a four-stranded antiparallel β-sheet with two α-helices packed against the β-sheet ^4^. Specific amino acids present in the four β-strands and in the loops connecting the secondary structure elements are responsible for the interaction of the RRM with, usually, single-stranded RNA. Despite the high conservation of the RRM fold, subtle amino acid differences have driven the evolution of RRMs towards the binding of highly different RNA sequences. Given that the RRM fold is the most abundant RNA binding domain, there are substantial variations in the binding specificity and interfaces of RRM folds to RNA ^5^, and even protein-protein interactions mediated by the helical region ^6,7^.

This suggests the possibility of rationally designing RRMs towards the selective, high-affinity, binding of specific RNA motifs, which can open novel strategies for the development of bio-analytical tools for in-cell RNA visualisation ^8–10^ or for the delivery of therapeutic RNAs protected in ribonucleoprotein (RNP) complexes similarly to Cas9/gRNA ^11,12^. The rational design of such protein mutants and modified RNA sequences requires the use of computational tools based on accurate structural information and an experimental characterization of the interaction by biophysical methodologies. In this respect, we have recently developed a computational tool that estimates and scores the binding preference between RRM domains and a given RNA sequence (RRMScorer) ^13^.

In this project we report the redesign of a tandem RRM protein and chose the Musashi protein given its biomedical importance and the fact that its RNA binding can be regulated by small molecule ligands ^14,15^. Overexpression of Musashi (MSI) proteins has been found in several malignant tumors. Importantly, there is a proposed correlation between the protein’s expression level, the proliferative activity of cancer cells, and a poor prognosis ^16,17^. There are two members of the mammalian MSI family: Musashi-1 (MSI-1) and Musashi-2 (MSI-2), which share a 69% sequence identity. MSI-1 contains an N-terminal disordered region followed by two tandem RRMs connected by a short inter-domain linker, about 10 amino acids long, and a large, disordered tail at the C-terminus. Both RRMs are involved in interactions with RNA while the C-terminal region is known to bind poly(A)-binding proteins ^17^ and has been associated with protein aggregation ^18^ and the formation of cellular condensates ^19^.

Human Musashi-1 is gradually down-regulated during neural differentiation and is involved in maintaining the undifferentiated state of neural stem cells through post-transcriptional control of downstream genes ^20,21^. Once activated, it regulates hundreds of different 3’-UTR regions of mRNAs and therefore it is implicated in various signalling pathways, including Notch and Wnt. Due to its regulatory functions, any alteration of the level of expression of Musashi-1 often leads to a disruption of signalling pathways, leading to several diseases, including cancer ^4^. This highlights the importance of MSI-1, and remarks on its potential to be a marker and a promising therapeutic target in cancer disease ^2,22,23^.

Musashi-1, as many other RBPs, has been considered for many years as an “undruggable” target due to the lack of a well-defined binding pocket. In recent years, several computational and experimental approaches have been developed to try to find small molecule inhibitors for proteins like Musashi-1. These approaches focus on trying to disrupt the RNA-protein interaction by blocking the binding interface, altering the protein structure, affecting its dynamics, varying its affinities ^15,24^ or degrade the upregulated protein ^25^. All these however, require a good understanding of the RNA-protein binding interface and mode of interaction. In case of Musashi-1 protein, structural information of mouse MSI-1 in complex with several short single-stranded RNA fragments ^26,27^ along with in vitro SELEX experiments ^28^ on the same protein have provided insight into the RNA binding specificity of each RRM domain of the protein. A consensus sequence (G/A)U_1-3_AGU has been reported to have a high affinity for the mouse MSI-1 RRM-1, while a preference for the generic UAG motif was observed for MSI-1 RRM-2 ^26^. However, there is no reported information for the human MSI-1, and more importantly, there is a lack of information on how the two RRMs interplay with RNAs of different lengths, compositions, and structures. Understanding the mode of binding of the tandem domain protein with different RNA could give valuable insights on the interaction and the binding interfaces implicated and could provide important structural information for the design of inhibitors.

Here, we have expressed in *E. coli* the two isolated domains (RRM-1 and RRM-2), and the tandem domain of human MSI-1 (RRM_1-2_), lacking the C-terminal tail (**Figure 1**). We designed protein mutants and altered RNA ligand sequence designed with RRMScorer to shed light on the binding mechanism between the protein and RNA. Specifically, we have employed complementary biophysical techniques to dissect and investigate the contributions to the interaction between the different constructs of the protein and selected RNA strands. For this we have combined NMR spectroscopy, Surface Plasmon Resonance (SPR), fluorescence quenching assays, and size-exclusion chromatography with multi-angle light scattering (SEC-MALS) to obtain comprehensive information on the binding sites, stoichiometry, binding kinetics, and affinity of the interactions. Our results reveal a competition between the two RRMs present in the tandem RRM_1-2_ when recognizing the same RNA sequence, leading to a more complex and dynamic interaction than expected. In this regard, the protein mutants and the RNA constructs designed by RRMScorer allowed us to investigate the contribution to the binding of the residues located in specific positions of the RRM domains, as well as that of nucleotides within the RNA strands. The results obtained using the mutated proteins and RNA constructs prove the validity of the strategy to use computational design for sequence specific RRM-RNA interactions.

**Figure 1.**
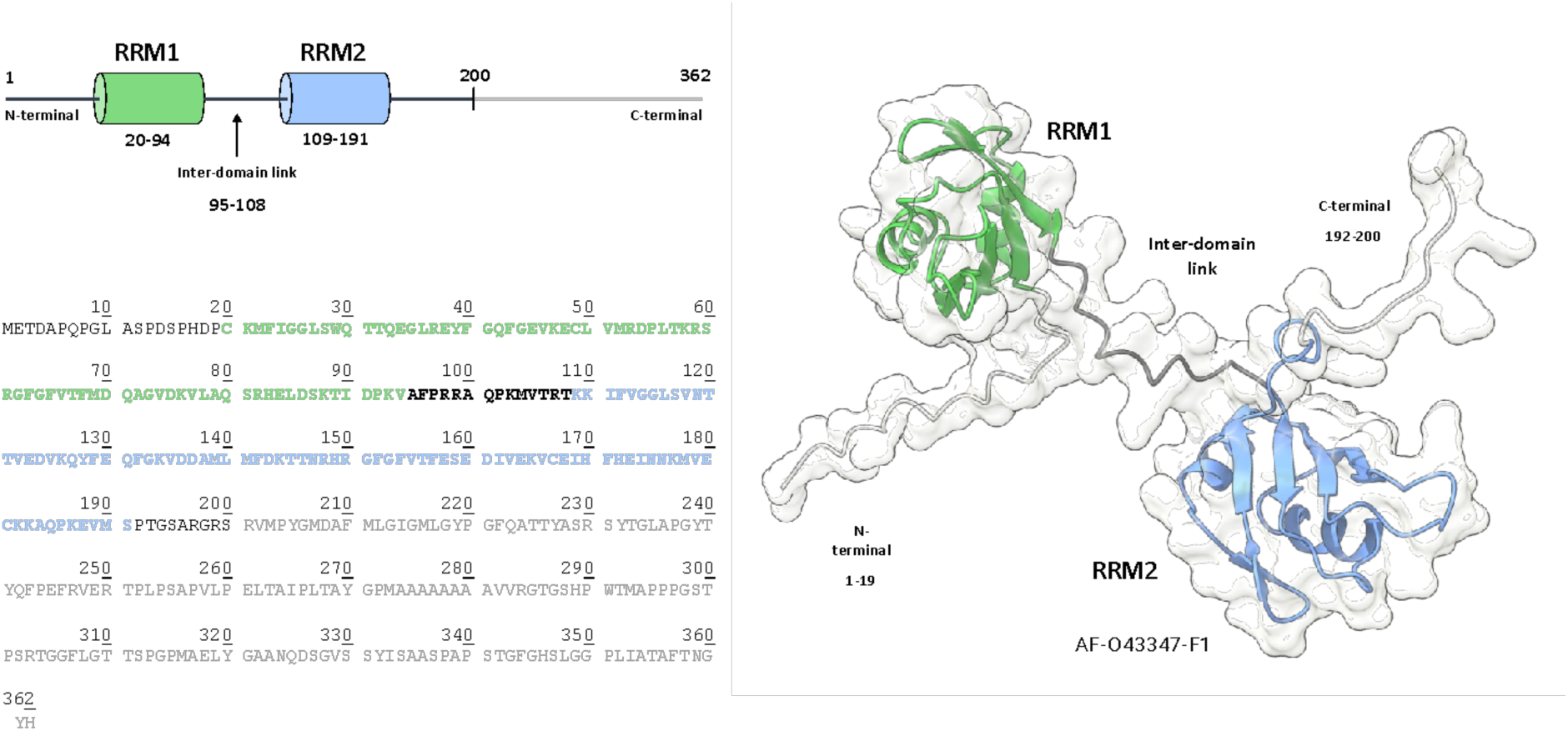
Schematic representation, sequence, and Alpha Fold model of human Musashi-1 RRM_1-2_ (1-200). Highlighted in green, black, and blue the RRM-1, inter domain linker, and RRM-2, respectively.

## Methods

### RNA strands

Synthetic **single stranded (L) and hairpin (HP) RNA** (**oligos 2-5** wild-type and mutants), for NMR experiments were purchased from Metabion international AG and Integrated DNA Technologies (IDT) (**Schematics and Table 1**). Biotinylated RNA oligonucleotides (**oligos 1-5** wild-type and mutants, **Schematics and Table 1**) used for SPR kinetic experiments were purchased from Metabion, Planegg, Germany. RNA used for fluorescence quenching assays (**oligo-HP4** and **oligo-HP5** constructs) conjugated with a 6-FAM fluorophore at the 5’-end and a BHQ-1 at the 3’-end were purchased from Integrated DNA Technologies (IDT).

**Schematics and Table 1.**
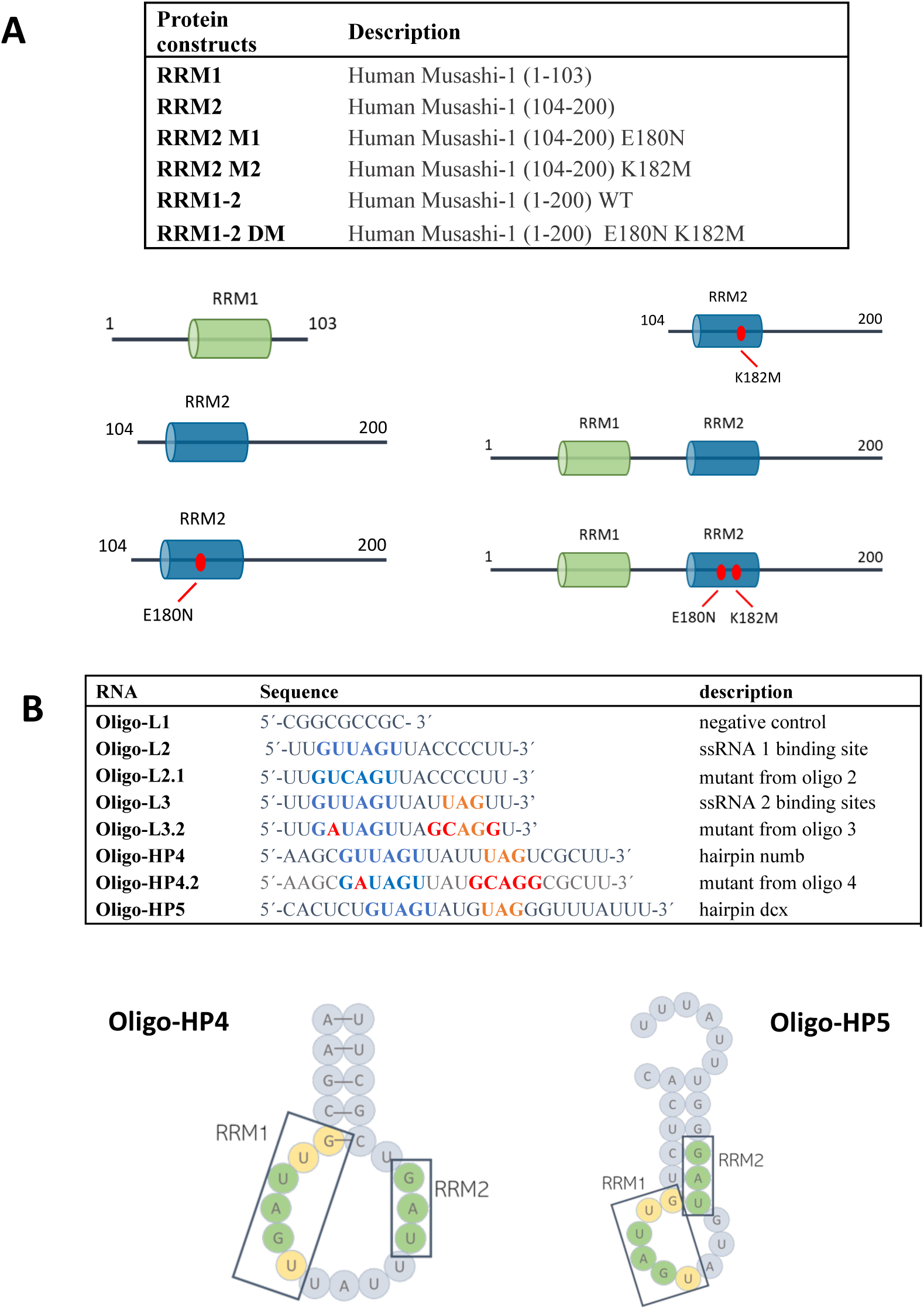
Description of A) protein constructs and B) RNAs used in this study. Highlighted the binding region for each RRM on the RNA sequences.

### Expression and purification of recombinant wild-type single RRM-1 and RRM-2 and tandem domains (RRM_1-2_) of human Musashi-1 (MSI-1)

Recombinant human Musashi-1 proteins (RRM-1: 1-103, RRM-2: 104-200 and RRM_1-2_: 1-200) were overexpressed in BL21(DE3) *E. coli* cells. Cells were grown in LB or M9 minimal media supplemented with ^15^NH_4_Cl or ^15^NH_4_Cl and ^13^C-glucose at 37 °C until optical density (OD600) reached 0.6-0.8. Expression was induced with 0.5 mM of isopropyl β-D-thiogalactoside (IPTG), cells were incubated at 37 °C for 3 h and harvested by centrifugation at 4 °C, for 15 min at 7500 rpm. Expression and purification steps for each construct are described in detail in the **Supplementary material**.

### Expression and purification of recombinant human Musashi-1 (MSI-1) RRM_1-2_ tandem domains E180N/K182M mutant

To produce the mutant, site-directed Mutagenesis was performed on the gene coding for the MSI-1 (1-200) RRM_1-2_ WT to replace the E180 by an asparagine and the K182 by a methionine to obtain the double mutant construct MSI-1 RRM_1-2_ E180N K182M (MSI-1 RRM_1-2_ DM). Recombinant human MSI-1 RRM_1-2_ DM protein in plasmid pET29b was overexpressed in BL21(DE3) GOLD *E. coli* cells. Expression was performed as described in the **Supplementary material** following the same protocol as for the wild-type tandem domain protein.

### NMR measurements and protein assignment

Experiments for the backbone resonance assignment (3D ^1^H-^15^N-^13^C HNCA, HNCACB and HNCO) were performed on ^13^C, ^15^N isotopically enriched samples of (MSI-1) RRM_1-2_ domain at the protein ^29^ concentration of 300 µM in buffer solution [20 mM MES pH 6.0, 100 mM NaCl, 1 mM DTT, 1 mM protease inhibitors]. NMR spectra were recorded at 298 K on a Bruker AvanceNEO NMR spectrometer operating at 1.2 GHz (^1^H Larmor frequency) and equipped with a TCI 3 mm cryo-probe.

Spectra were processed with the Bruker TOPSPIN software packages and analysed with CARA (Computer Aided Resonance Assignment, ETH Zurich). The backbone resonance assignment of MSI-1 RRM_1-2_ was obtained by comparing the assignments available in the literature for the individual domains from the mouse MSI-1 protein (BMRB code: 11450 and 36058) ^26,27^ with the NMR spectra recorded on MSI-1 RRM_1-2_ and analysing triple resonance spectra recorded on MSI-1 RRM_1-2_.

### R1, R2 and NOE measurements

For the characterization of human Musashi-1 RRM_1-2_ tandem domain protein, NMR measurements, R1, R2 and NOE measures and protein assignment have been performed on the ^15^N-enriched sample and can be found in the **Supplementary Materials** ^29^.

### Computational design of protein and RNA mutants

For the design of protein mutations and new RNA constructs, scoring of RRM-RNA interactions upon mutations in both Musashi-1 and RNA targets were computed with the RRMScorer method ^10^ (https://bio2byte.be/rrmscorer/). RRMScorer estimates the binding preferences between any specific RRM and a given RNA solely based on their sequences. In summary, from all available structures available in PDB for the RRM-RNA complexes a set of entries, describing the canonical binding mode of RRM domains, ^1^ was selected after a careful alignment of the RRM-RNA complexes. The contacts observed within this set of structures, and the positions of the residues involved in the binding on the RRM domains, were integrated in a probabilistic framework to extract propensities for residue-nucleotide contact preferences in specific positions. This method appeared to be particularly suitable for the limited amount of data and residue-level information that is currently available. The most relevant positions of the protein regarding RNA binding are short-listed based on RRMScorer database contacts analysis, and for each of them, a preference matrix has been derived showing the binding preferences to a nucleotide for any residue in that specific position.

### Titration of Musashi with RNA strands

The effect of linear (L) single stranded RNA constructs **oligo-L2**, **oligo-L3,** and **oligo-L3.2** (**Schematics and Table 1**) was evaluated on ^15^N-isotopically enriched MSI-1 RRM_1-2_, MSI-1 RRM-1 and MSI-1 RRM-2 proteins at the concentration of 100 μM in the following experimental conditions: 50 mM Tris-HCl, 140 mM NaCl, 1 mM EDTA, 1 mM protease Inhibitors. The interaction with three other RNA strands **oligo-HP4**, **oligo-HP4.2,** and **oligo-HP5** (**Schematics and Table 1**) able to form a hairpin (HP), was also investigated under the same experimental conditions. The pH was 7.2 in the case of MSI-1 RRM_1-2_ and MSI-1 RRM-2, while the pH was 7.5 in the case of MSI-1 RRM-1, taking into account its lower isoelectric point. The effect of RNA constructs (**oligo-L2, oligo-L3, oligo-L3.2, oligo-HP4** and **oligo-HP4.2**) (**Schematics and Table 1**) was also evaluated on ^15^N-isotopically enriched MSI-1 RRM_1-2_ DM protein at the concentration of 100 μM in the following experimental conditions: 50 mM Tris-HCl, 140 mM NaCl, 1 mM EDTA, 1 mM protease Inhibitors. 2D ^1^H ^15^N HSQC and 2D ^1^H ^15^N TROSY NMR spectra were recorded at 298 K on the single domains and on the tandem domain, respectively, using a Bruker AvanceNEO NMR spectrometer, operating at 900 MHz (^1^H Larmor frequency). During the NMR titrations, increasing amounts of the RNA strands were added to the protein solution to reach the final concentrations of 6, 12, 24, 50, 120, 200 μM of RNA. Spectra were processed with the Bruker TOPSPIN software packages and analysed with CARA (Computer Aided Resonance Assignment, ETH Zurich).

### Size-Exclusion Chromatography with Multi-Angle Light scattering (SEC-MALS)

Samples of 100 µL were loaded at 0.6 mL/min on a Superdex 200 10/300 GL analytical size-exclusion column (GE Healthcare), and elution was monitored by the following in-line detectors: a light scattering diode array (DAWN EOS, Wyatt Technology UK Ltd.), a dynamic module (WYATT QELS, Wyatt Technology UK Ltd.), UV detector (Smartline UV Detector 2500, Knauer) and a differential refractive index detector (Optilab rEX,Wyatt Technology UK Ltd.).

Chromatograms were analysed using the ASTRA software (v7.3.2.19, Wyatt Technology UK Ltd.) and the interaction chromatograms were analysed using the Protein Conjugate template.

Parameters of the specific refractive index increment dn/dc (mL/g) and UV Extinction coefficient (mL/(mg·cm)) of each domain (RRM-1 and RRM-2) and of the modifiers (**oligo-L2**, **oligo-L3**, **oligo-HP4** and **oligo-HP5**) are found in **Table S1**.

### MSI-1-RNA interaction kinetics experiments

Kinetic experiments were performed using Surface Plasmon Resonance (SPR) based biosensor Biacore 3000 and T 200 (Cytiva). Data analysis was performed using TraceDrawer 1.9.2 and 1.10 (Ridgeview Instruments). A more complete experimental methodology and data analysis are described in the **Supplementary material**.

### Fluorescence quenching assays

The RNA **oligo-HP4** and **oligo-HP5** constructs conjugated with a 6-FAM fluorophore at the 5’-end and a Black Hole Quencher 1 (BHQ1) ^18^ at the 3’-end were purchased from Integrated DNA Technologies (IDT). The oligos were resuspended with NMR buffer (50 mM Tris pH 7.2, 140 mM NaCl and 1 mM EDTA) and snap-cooled by heating at 95°C for 5 min followed by incubation in ice for 15 min. All samples were diluted to a final RNA concentration of 400 nM and MSI-1 added to reach the desired molar ratio. A denaturing RNA control for both **oligo-HP4** and **oligo-HP5** were prepared by adding 8 M urea and heating the sample at 95 °C before the measurement. FAM fluorophore was excited at 480 nm and emission recorded at 520 nm. The standard deviation was determined performing three replicates. Formation and unfolding of the RNA hairpins upon MSI-1 titration is monitored based on the fluorescence emission intensity compared to a denaturing control (**Suppl. Fig. 9**) and an RNA-only control.

### NMR spectroscopy for quenching assays

NMR samples were prepared from stock RNA samples after snap-cooling by heating at 95 °C for 5 min and incubation in ice for 15 min. 1D ^1^H experiments were collected on samples of RNA at the concentration of 50 μM in NMR buffer (50 MM Tris pH 7.2, 140 mM NaCl and 1 mM EDTA) with 10% D_2_O, added as lock signal and acquired at 298 K on 800 MHz Bruker Avance NMR spectrometer, equipped with cryogenic triple resonance gradient probe. NMR spectra were processed with TopSpin 3.5.

## Results

### NMR characterization of free MSI-1

A set of 2D ^1^H-^15^N -HSQC spectra of the three constructs (MSI RRM_1-2_ tandem domain, RRM-1, and RRM-2 isolated domains), was recorded and superimposed to evaluate the folding state and interdomain flexibility of the tandem RRMs of human MSI-1 (see **Figure 2A**). All spectra show well-dispersed signals in agreement with folded protein structures. The NMR spectrum of the MSI RRM_1-2_ tandem domain is largely superimposable to the spectra of the isolated domains, as the majority of signals in the spectrum of MSI RRM_1-2_ overlap with either RRM-1 or RRM-2 signals. The absence of a large chemical shift perturbations when comparing the isolated domains with the tandem construct suggests a large interdomain flexibility with few interactions between the two domains or with the linker in the tandem domain construct. In the MSI RRM_1-2_ spectrum, several additional signals, that can be attributed to the portion of the interdomain linker, are also visible. NMR ^15^N relaxation data (**Figure 3**) corroborate these findings by showing the presence of a sizable interdomain flexibility. The backbone assignment of the MSI RRM_1-2_ tandem domain was obtained starting from the published assignments of the murine isolated RRM domains (BMRB codes: 11450 and 36058) ^26,27^ considering the high sequence homology between the mouse and human MSI-1 proteins (99.44%). This was confirmed and complemented by the analysis of triple resonance spectra (see **Supporting Information**). 92.5% of all residues, including those forming the linker region, have been assigned (**Figure 2B**) and the protein resonance assignment has been reported in the Biological Magnetic Resonance Data Bank (BMRB) under the accession code: 52590.

**Figure 2.**
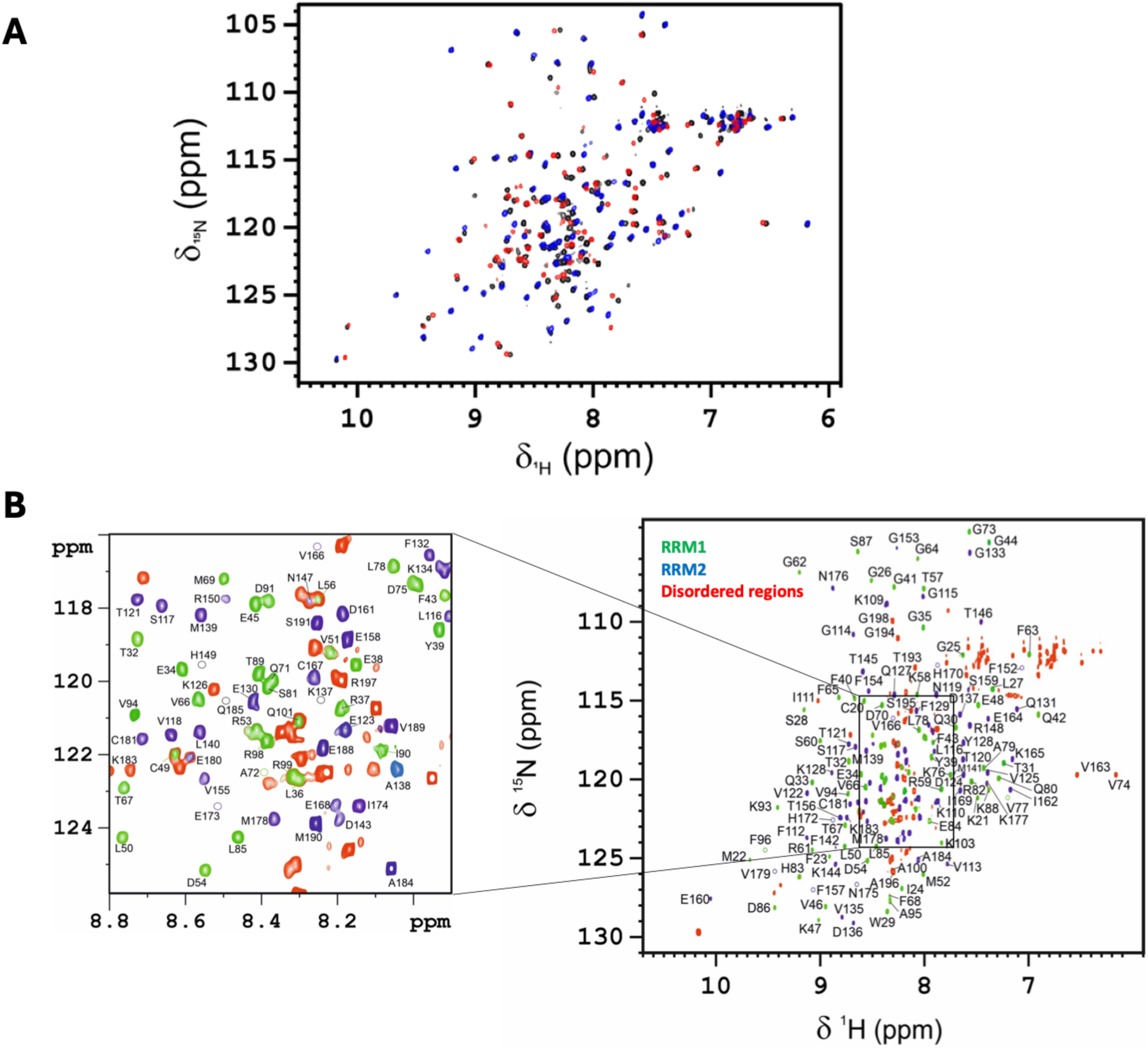
(A) Superimposed 2D ^1^H-^15^N HSQC spectra of isolated RRM-1 and RRM-2, and TROSY spectrum of MSI-1 RRM_1-2_ tandem domain. (B) Assignment in a 2D ^1^H–^15^N HSQC of the ^13^C, ^15^N isotopically enriched (MSI-1) RRM_1-2_ domain at 900 MHz and 298 K. Highlighted in green, blue and red the RRM-1, RRM-2 and disordered regions, respectively.

**Figure 3.**
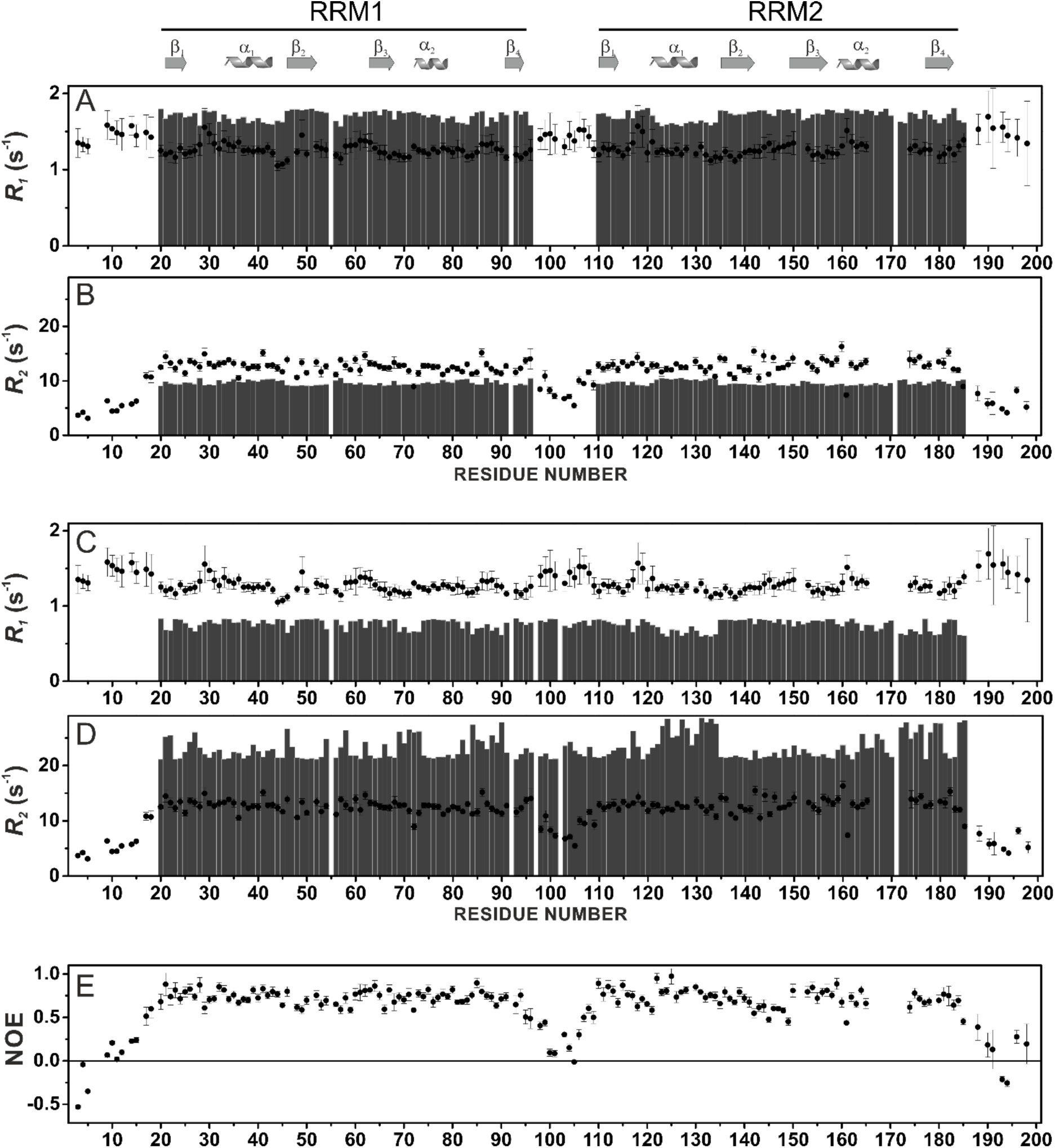
Comparison of experimental backbone ^15^N_H_ R_1_ values for MSI-1 RRM_1-2_ construct (data collected at 298 K, black filled circles) with the calculated values (grey bars) for isolated RRM-1 and RRM-2 domains (A) and for the full MSI-1 RRM_1-2_ tandem domain construct (C). Comparison of experimental backbone ^15^N_H_ R_2_ values for MSI-1 RRM_1-2_ construct (data collected at 298 K, black filled circles) with the calculated values (grey bars) for isolated RRM-1 and RRM-2 domains (B) and for the full MSI-1 RRM_1-2_ tandem domain construct (D). Experimental NOE values for MSI-1 RRM_1-2_ construct (data collected at 298 K) (E).

### RRM domains compete for the recognition of the (A/G)U_1-3_AGU motif

While RNA binding studies have been reported and binding motifs for the RRMs have been reported ^17,20,26,27^, the binding mechanism of the human Musashi-1 is still largely unexplored. Considering the very high sequence homology with the better-characterised mouse protein, it is likely that the minimal RNA sequence recognized by RRM-1 and RRM-2 domains is UAG. In addition, an *in vitro* selection for high-affinity RNA ligands for MSI-1 identified a longer consensus recognition sequence (G/A)U_1-3_AGU for RRM-1 ^26–28^. Therefore a linear (L) RNA oligonucleotide (**oligo-L2:** 5’-UU**GUUAGU**UACCCCUU-3’) bearing a single consensus binding motif (G/A)U_1-3_AGU was designed to explore the binding mechanism of MSI-1.

The interaction of MSI-1 with **oligo-L2** was investigated by recording 2D ^1^H-^15^N -HSQC NMR spectra of the three protein constructs (MSI RRM_1-2_ tandem domain, RRM-1 and RRM-2 isolated domains) before and after the addition of increasing amounts of the RNA strand. The interactions between RRM-1 and RRM-2 with **oligo-L2** are in the slow exchange regime on the NMR timescale (**Figure 4**) in agreement with a high affinity of the oligonucleotide for the two proteins. SEC-MALS and kinetic SPR experiments confirmed a 1:1 binding model, with dissociation constants (K_D_) (**Figure S5, S7, Figure 8 and Table 2**) in the nanomolar range (see **Suppl. Information**). Since the UAG sequence is included in the (G/AU_1-3_AGU) motif, the high affinity of RRM-2 for **oligo-L2** is not unexpected. Conversely, the analysis of the interaction between the MSI-1 RRM_1-2_ tandem domain and **oligo-L2** was somewhat surprising. When the tandem domains are titrated with increasing amounts of **oligo-L2**, some cross-peaks of the free protein broaden and decrease in intensity (**Figure 4**) with some signals experiencing small Chemical Shift Perturbations (CSP) (**Figure S2**). At the same time, cross-peaks of a new species could not be clearly detected, and even with an excess of RNA with respect to the protein (protein/RNA molar ratio of 1:2), we could hardly see new signals, and those present had very low intensity. Interestingly, the signals of residues belonging to the β-sheet surface of both domains are affected after the addition of **oligo-L2** (**Figure S2**). This behaviour suggests a competition between the two domains for the binding of **oligo-L2**. The line-broadening observed in the NMR titrations is also consistent with this, where alternating binding in fast to intermediate exchange to the two RRMs with varying chemical environments can rationalise the line-broadening. This is further confirmed by the results of kinetic studies as presented in **Figure 8** and **Table 2**. Kinetics of this interaction were evaluated with SPR and the obtained sensorgrams for the isolated RRM-1 and RRM-2 domains are very similar (**Figure S7**) and showed interactions with fast association and dissociation rates. For the MSI RRM_1-2_ tandem domain, a similar fast association was seen (**Figure S7**), but it was followed by a biphasic dissociation, which is initially fast but rapidly slows down, suggesting a stabilisation of the interaction. This behaviour has been already attributed to the formation of bivalent interactions between the tandem domain and the RNA, when the amount of unbound RNA is sufficient ^30^. The calculated affinities (K_D_) to RRM_1-2_ were in the nanomolar range for the first monovalent interaction and in the lower micromolar range for the bivalent interaction (see **Figure 8, Table 2** and **Suppl. Material**).

**Figure 4.**
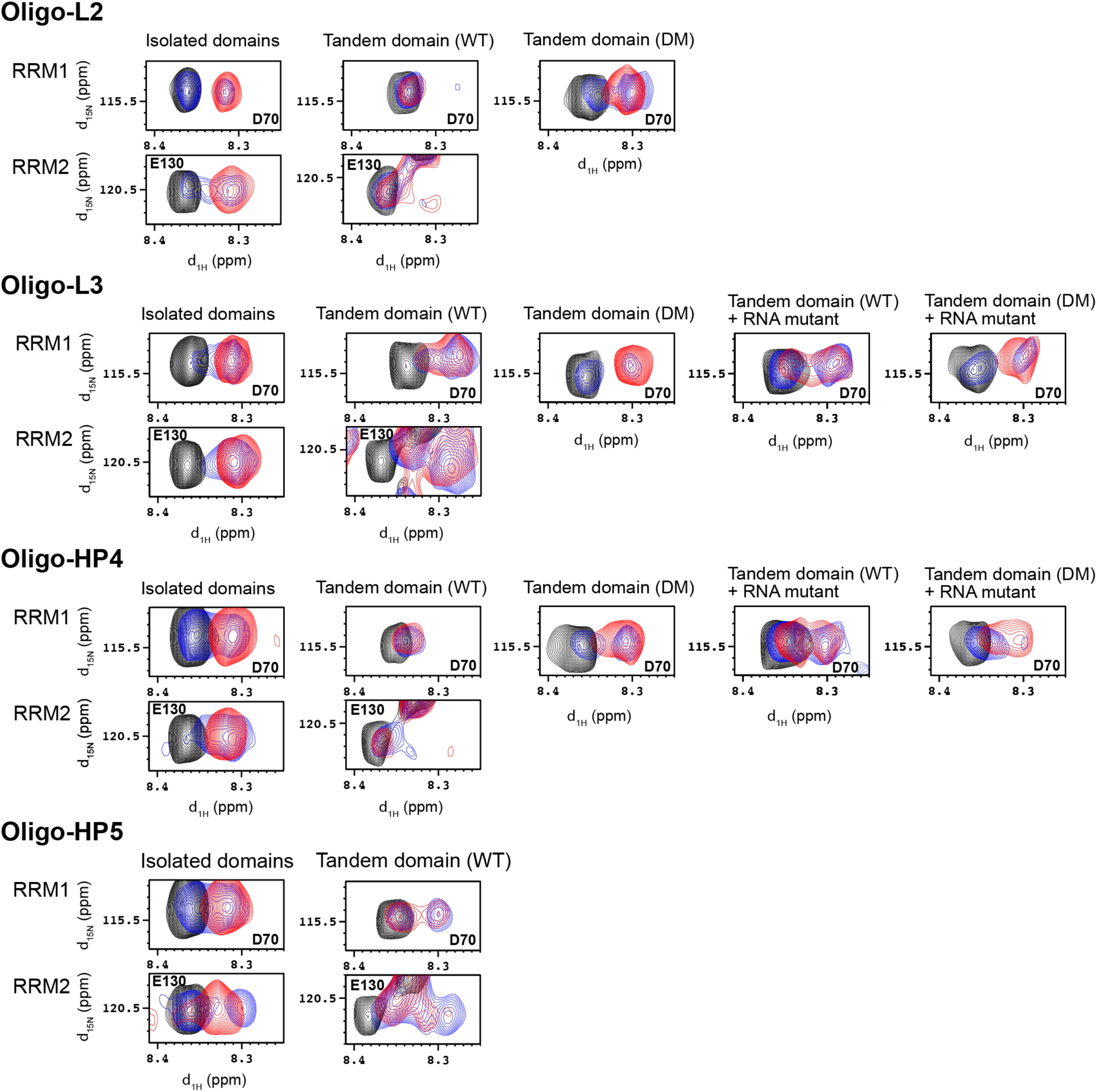
Zoom views of the 2D ^1^H-^15^N HSQC and TROSY spectra, recorded on the isolated RRM-1 and RRM-2 domains, on the tandem domain wild-type (WT) and double mutant (DM: E180N, K182M), respectively. In black are the spectra of the free proteins, in blue the spectra of the proteins in the presence of sub-stoichiometric concentrations of **oligo-L2**, **oligo-L3**, **oligo-L3.2**, **oligo-HP4**, **oligo-HP4.2** or **oligo-HP5** (protein/RNA ratio of about 1:0.5), and in red the spectra of the proteins in the presence of **oligo-L2**, **oligo-L3**, **oligo-L3.2**, **oligo-HP4**, **oligo-HP4.2** or **oligo-HP5** in the protein/RNA ratio of 1:1. The signals assigned to Asp-70 and Glu-130 are displayed in the figure for RRM-1 and RRM-2 domain, respectively.

**Table 2.** Kinetic and affinity mean values of isolated domains with **oligos 2 – 5** based of triplicates.

|  | *MSI-1 RRM-1* |  |  | *MSI-1 RRM-2* |  |  |
| --- | --- | --- | --- | --- | --- | --- |
| | $k_{a1}$ ($M^{-1}s^{-1}$) | $k_{d1}$ ($s^{-1}$) | $K_{D1}$ (nM) | $k_{a1}$ ($M^{-1}s^{-1}$) | $k_{d1}$ ($s^{-1}$) | $K_{D1}$ (nM) |
| *Oligo-L2* | $(3.5 \pm 0.2) 10^6$ | $0.167 \pm 0.010$ | $47.1 \pm 0.7$ | $(4.6 \pm 0.7) 10^6$ | $0.112 \pm 0.007$ | $25.0 \pm 3.6$ |
| *Oligo-L2.1* | $(2.6 \pm 0.9) 10^6$ | $0.370 \pm 0.062$ | $151 \pm 16.9$ | $(1.8 \pm 0.2) 10^6$ | $0.218 \pm 0.008$ | $123 \pm 7.8$ |
| *Oligo-L3* | $(4.5 \pm 0.1) 10^5$ | $0.069 \pm 0.003$ | $153 \pm 8.6$ | $(4.7 \pm 0.2) 10^5$ | $0.038 \pm 0.0004$ | $81.6 \pm 2.0$ |
| *Oligo-L3.2* | $(1.4 \pm 0.3) 10^6$ | $0.120 \pm 0.032$ | $85.6 \pm 7.4$ | $(2.8 \pm 1.2) 10^6$ | $0.121 \pm 0.047$ | $44.5 \pm 4.5$ |
| *Oligo-HP4* | $(8.3 \pm 0.4) 10^6$ | $0.178 \pm 0.013$ | $21.9 \pm 2.7$ | $(1.1 \pm 0.1) 10^6$ | $0.038 \pm 0.004$ | $37.2 \pm 2.4$ |
| *Oligo-HP5* | $(8.0 \pm 0.5) 10^6$ | $0.268 \pm 0.051$ | $32.8 \pm 5.2$ | $(2.1 \pm 0.1) 10^6$ | $0.055 \pm 0.002$ | $26.4 \pm 1.6$ |
|  | *MSI-1 RRM-2 E180N (RRM2-M1)* |  |  | *MSI-1 RRM-2 K182M (RRM2-M2)* |  |  |
| | $k_{a1}$ ($M^{-1}s^{-1}$) | $k_{d1}$ ($s^{-1}$) | $K_{D1}$ (nM) | $k_{a1}$ ($M^{-1}s^{-1}$) | $k_{d1}$ ($s^{-1}$) | $K_{D1}$ (nM) |
| *Oligo-L2* | $(8.4 \pm 1.1) 10^6$ | $0.288 \pm 0.015$ | $35.5 \pm 3.6$ | $(9.6 \pm 1.6) 10^5$ | $0.100 \pm 0.01$ | $108 \pm 10.2$ |
| *Oligo-L2.1* | $(6.7 \pm 0.9) 10^6$ | $0.389 \pm 0.044$ | $58.6 \pm 3.6$ | $(3.5 \pm 0.2) 10^6$ | $0.227 \pm 0.05$ | $62.9 \pm 11.0$ |
| *Oligo-L3* | $(7.0 \pm 0.9) 10^6$ | $0.507 \pm 0.003$ | $74.8 \pm 9.3$ | $(1.5 \pm 0.1) 10^6$ | $0.094 \pm 0.002$ | $64.5 \pm 4.1$ |
| *Oligo-L3.2* | $(4.2 \pm 1.1) 10^6$ | $0.187 \pm 0.025$ | $49.8 \pm 11.4$ | $(1.0 \pm 0.05) 10^6$ | $0.187 \pm 0.005$ | $88.5 \pm 1.25$ |
| *Oligo-HP4* | $(1.4 \pm 0.3) 10^7$ | $0.178 \pm 0.011$ | $14.1 \pm 2.5$ | $(1.1 \pm 0.7) 10^7$ | $0.177 \pm 0.035$ | $39.1 \pm 15$ |
| *Oligo-HP5* | $(2.6 \pm 0.3) 10^7$ | $0.375 \pm 0.036$ | $14.5 \pm 0.8$ | $(7.7 \pm 0.5) 10^6$ | $0.112 \pm 0.026$ | $14.5 \pm 3.0$ |

Following our observation of this competing phenomenon, we investigated how the two domains, both separately and in tandem, behave when two binding sites are present simultaneously in a linear RNA strand. Therefore, a new RNA oligonucleotide (**oligo-L3:** 5’-UU**GUUAGU**UAU**UAG**UU-3’) containing the consensus sequences known to bind both RRM-1 and RRM-2 domains (G/A)U_1-3_AG) ^28^ and UAG motifs ^26^, respectively, was designed. Firstly, the two isolated RRM domains were titrated separately with **oligo-L3** and the interaction monitored by NMR. As observed for **oligo-L2**, the interactions of isolated RRM-1 and RRM-2 with **oligo-L3** are in the slow exchange regime on the NMR timescale (**Figure 4**). The data indicate that RRM-1 and RRM-2 can bind both RNA motifs, with the formation of complexes with different protein/RNA stoichiometric ratios as described in detail in the **Supplementary Material** (**Figure S5**). The NMR data are supported by SEC-MALS analysis, which was carried out to shed light on the stoichiometry of the interaction between RRM-1 and **oligo-L3**. SEC-MALS chromatogram analysis confirmed the presence of two species in solution, the most abundant one with a 2:1 protein/RNA stoichiometry, and a minor one with a 1:1 protein/RNA stoichiometry (see **Figure S5 and Suppl. Materials**). Kinetics for this interaction also show high-affinity binding within the nanomolar range with an almost 8-fold slower association and less than 3-fold slower dissociation rate constants when comparing with **oligo-L2** (**Figure 8**, **Table 2 and Supplementary Material**). Altogether these results highlight once again this competitive phenomenon.

Secondly, we investigated the tandem domain protein interacts with **oligo-L3**. Solution NMR showed a decrease in the intensity of the cross-peaks of the free protein upon the addition of **oligo-L3**. New cross-peaks, corresponding to the MSI-1 RRM_1-2_ tandem domain in complex with **oligo-L3**, appear and increase in intensity as well (**Figure 4**). Therefore, the interaction of the tandem domain with **oligo-L3** occurs in a slow exchange regime on the NMR timescale, unlike the observed interaction with **oligo-L2** (**Figure 4**). However, in the presence of RNA at a concentration of 100 µM (1:1 protein/RNA molar ratio), the new signals are still broad, so the absence of multiple conformational states cannot be excluded. Assignment of the newly shifted signals is not feasible because of the higher ambiguity due to the higher crowding in the spectra of the MSI-1 RRM_1-2_ tandem domain.

The kinetics of the interaction between the MSI-1 RRM_1-2_ tandem domain and **oligo-L3** showed a much slower association rate constant resulting in a 4-fold weaker affinity compared to **oligo-L2** (**Figure 8 and Table 3**). This reduction in the association rate, that was also observed for the isolated RRM domains, was unexpected and InteractionMap analysis was performed to further evaluate this behaviour. InteractionMap showed that the peaks corresponding to the mono- and bivalent interaction were similar for oligos containing the UAG motif (**oligo-L2, -L3, -HP4,** and **-HP5**) but for **oligo-L3** these peaks are broader with respect to the association rate constant indicating heterogeneous recognition (**Figure 7A**). As NMR results suggests that both UAG motifs can be bound by RRM domains, it is most likely that the first RRM that binds to **oligo-L3** results in a slower binding of a second RRM domain to the second UAG motif.

**Table 3.**
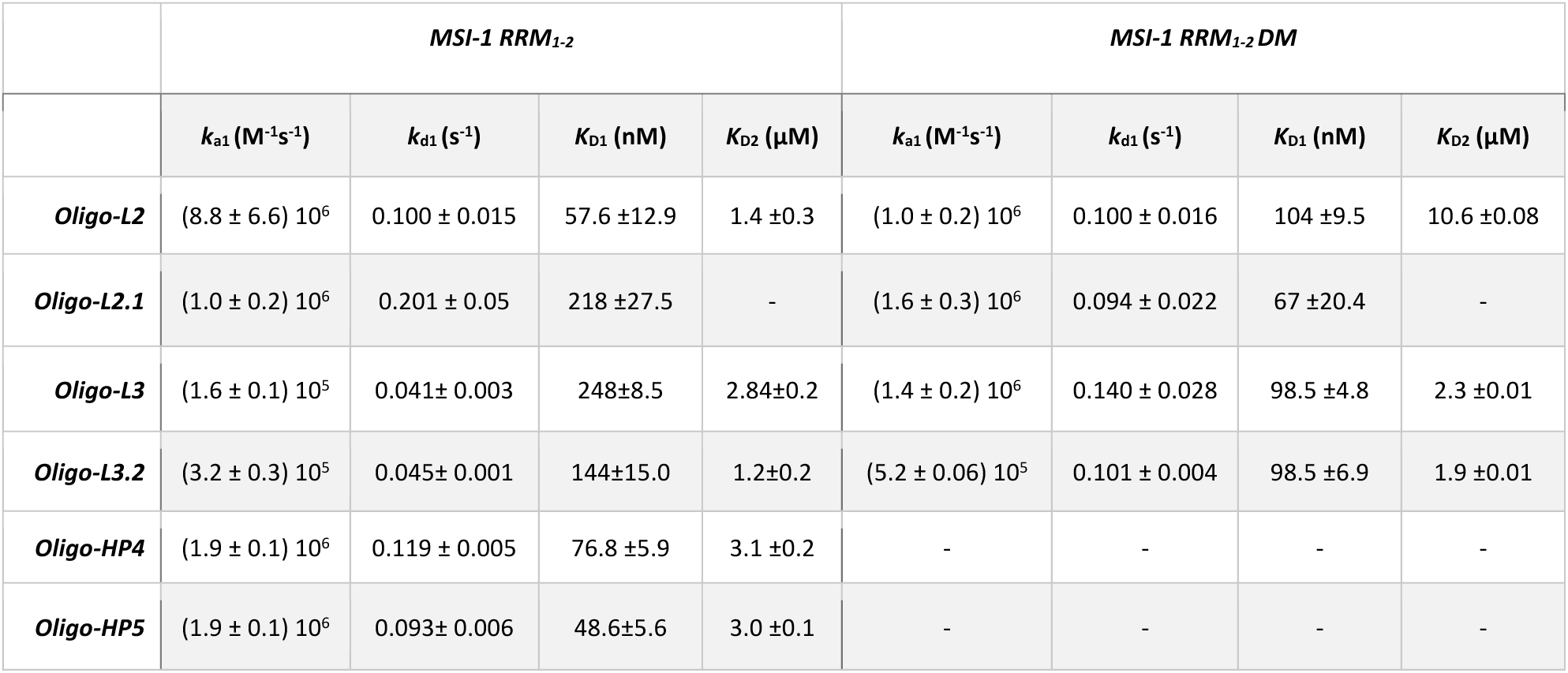
Kinetic and affinity mean values of tandem domain protein (MSI-1 RRM_1-2_ and MSI-1 RRM_1-2_ DM) with **oligos 2 – 5** based on triplicates.

### How secondary structures of RNA affect the binding with human Musashi-1 protein

Binding of Musashi-1 to folded RNA strands (i.e., hairpins (HP)) was then investigated using **oligo-HP4**(5’-AAGC**GUUAGU**UAUU**UAG**UCGCUU-3’) and **oligo-HP5** (5’-CACUCU**GUAGU**AUG**UAG**GGUUUAUUU-3’) that were selected because of the different locations of the consensus motifs, known to bind RRM-1 and RRM-2, respectively. **Oligo-HP4** is an RNA fragment from the *NUMB* mRNA ^28^ that contains both the binding motifs (the G/AU_1-3_AGU motif for RRM-1 and the UAG motif for RRM-2) within the loop region of a hairpin folding. **Oligo-HP5** is an RNA fragment from the *DOUBLECORTIN* (dcx) mRNA ^31^ and contains one consensus motif (G/AU_1-3_AGU) in the loop region, and the second UAG motif in the double-stranded region of the hairpin.

Solution NMR spectra show that in the presence of the hairpin constructs, the isolated RRM domains behave similarly to what is observed with the linear RNA sequence containing two binding sites (**oligo-L3,** see the previous section and **Supplementary material**). However, when the MSI-1 RRM_1-2_ tandem domain is titrated with the hairpin constructs, different behaviours are observed. In particular, in the NMR titration of **oligo-HP4** we observe only a decrease in signal intensity, without the appearance of new cross-peaks in the NMR spectra (**Figure 4**). Conversely, when the MSI-1 RRM_1-2_ tandem domain is titrated with sub-stoichiometric concentrations of **oligo-HP5**, new cross-peaks corresponding to the protein in complex with RNA are observed (**Figure 4**). Interestingly, also after the addition of an excess of the oligonucleotide with respect to the protein (∼ 200 μM, protein/RNA ratio of 1:2) the signals of the new species do not increase in intensity, and the signals of the free protein are still present in the spectrum. In this regard, competition between the two domains for the same RNA site, and the presence of multiple species in solution can be hypothesised. Furthermore, the interaction landscape may be complicated by the possibility of an opening of the hairpin structure (see **Suppl. Material**). Therefore, NMR data can give information on the binding regions, but not about the strength of the interaction.

To verify whether the RNA hairpins are destabilised by binding of MSI-1, we performed fluorescence quenching assays using **oligo-HP4** and **oligo-HP5.** Quenching of the fluorescence is seen for the hairpin as the fluorescent dye (6-FAM) and the quencher (BHQ1) are in close spatial proximity (**Figure 5, and Suppl. Material**) ^32,33^. The fluorescence quenching assay for the RNA-only and denaturing control samples shows a significant difference (**Figure S9**). Upon titration of the two oligos with MSI-1 RRM_1-2_, a concentration-dependent increase in the fluorescence intensity is observed compared to the control, consistent with the binding of MSI-1 to the single-stranded form of the RNA. This is further confirmed by ^1^H imino NMR spectra monitored upon titration of **oligo-HP4** and **oligo-HP5** with MSI-1 RRM_1-2_. Reduction of the intensity of the imino signals upon the addition of increasing amounts of MSI-1 RRM_1-2_ (**Figure 5B**) indicates unfolding of the RNA hairpin upon binding. Taken together, the fluorescence quenching assays, and NMR measurements indicate that both **oligo-HP4** and **oligo-HP5** adopt a hairpin conformation, that upon MSI-1 RRM_1-2_ binding is unfolded, leading the protein to interact with a single-stranded RNA conformation.

**Figure 5.**
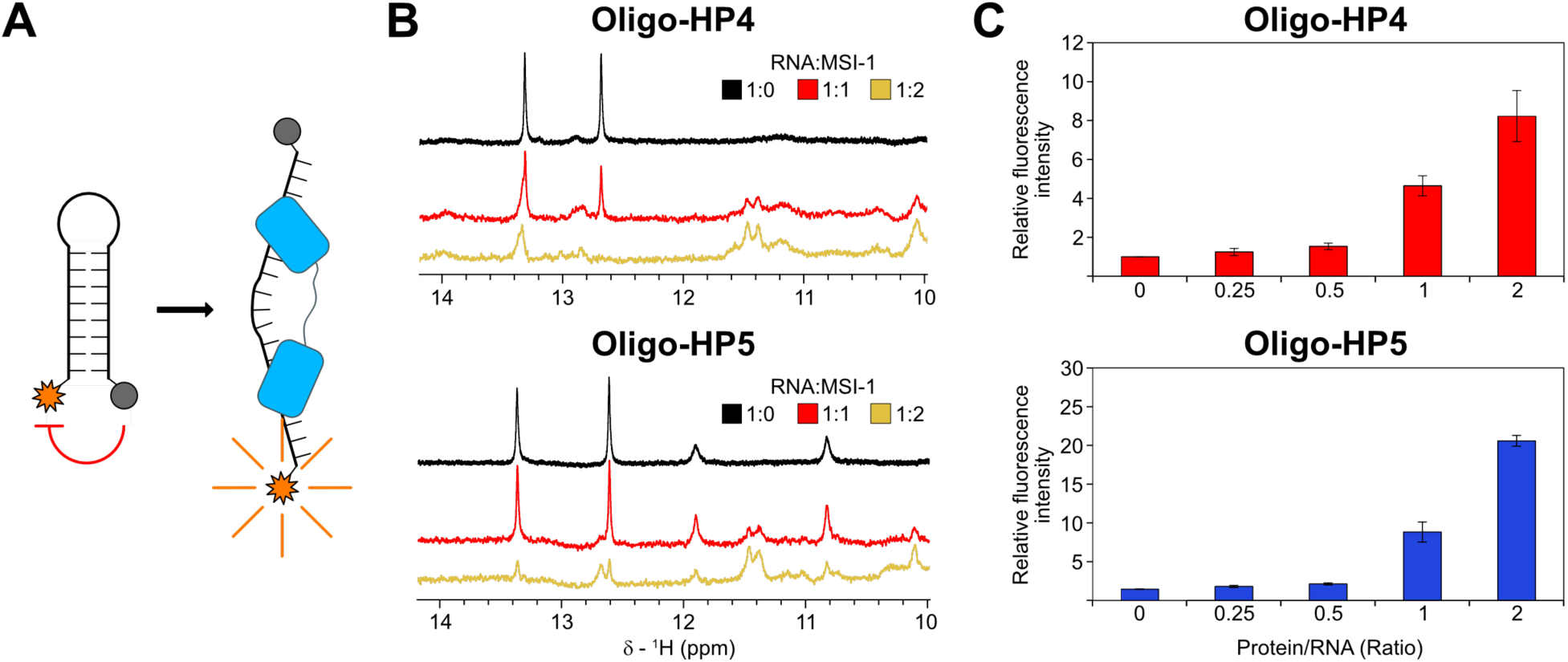
MSI-1 binding to **oligo-HP4** and **oligo-HP5** involves unfolding of their RNA hairpin structures. (A) Schematic representation of the label RNAs used for fluorescence quenching assays highlighting the conjugated fluorophore at the 5’ (in orange) and the quencher at the 3’ (in grey). When the two probe are not close in space (i.e. upon unwinding of the hairpin by protein binding), there is fluorescence emission by the fluorophore. (B) Monitoring of **oligo-HP4** and **oligo-HP5** upon addition of MSI-1 with imino NMR spectra and (C) fluorescence quenching assays (bar plots). For fluorescence quenching assays, fluorescence intensity values are normalized respect to the emission of the zero point of the titrations

Kinetics studies showed a similar binding of MSI-1 RRM_1-2_ tandem domain to both **oligo-HP4** and **oligo-HP5**, resulting in comparable kinetic rate constants and affinity values for both the monovalent and bivalent interactions (**Figure 8 and Table 2**). The affinities for the monovalent interactions are also similar to the values obtained for **oligo-L2**, while the affinities for the bivalent interactions for both **oligo-HP4** and **oligo-HP5** are similar to the affinity of the bivalent interactions obtained for **oligo-L3**. This confirms that the initial interaction (monovalent) of MSI-1 RRM_1-2_ with hairpins structures occurs in the same way as the interaction observed with the linear oligo bearing a single motif (**oligo-L2**), while the second interaction (bivalent) has a binding strength similar to that observed for the interaction with the linear oligo containing two binding motives (**oligo-L3)**.

### Design of protein mutants and RNA binding sites modifications to increase the affinity and specificity of protein-RNA interactions

To investigate the molecular bases of the competition between the two RRM domains for the same binding site, the preference of RRM-1 and RRM-2 for different RNA constructs was investigated using RRMScorer. First, the interaction of RRM-1 and RRM-2, respectively, with the 3-nucleotides sequence N_x_N_x_N_x_ (where N_x_ can be any nucleic base) was analysed with RRMScorer (**Figure 6**). As expected, a clear preference of both RRMs towards the UAG motif was observed, but a better score for affinity (0.1 difference in a logarithmic scale) was obtained for RRM-1. Second, the best possible scores of each RRM towards the 5-nucleotide motif N_x_UAGN_x,_ were evaluated again by RRMScorer. The best scores are highlighted in **Figure 6**, while the positions with higher scores and differences between both RRMs are plotted in **Figure S10**. The analysis identified several RNA motifs predicted to have a reduced affinity (lower score) for RRM-1, while keeping a relatively high score towards RRM-2 (**Figure 6**). An RNA construct with two different short motifs to specifically target RRM-1 and RRM-2, respectively, is expected to show a stronger affinity for the tandem domain, as it avoids the binding to both RRMs. In this regard, we aimed to design a new RNA that would contain a 3 to 5 nucleotide motif that would have lower affinity for RRM-1 and higher affinity for RRM-2 to try to redirect the preference of binding and understand the molecular bases of this competition. The CAG motif was identified as the most promising candidate for RRM-2, with CCAGG, and GCAGG the top-scoring 5-mers, to replace the UAG binding site in newly designed RNA strands (see **supplementary Figure S10**). Improvement in the specificity and affinity of binding can also be obtained by designing protein mutants with RRMScorer, using an almost identical approach and a mutant of the RRM-2 domain was designed with a decreased affinity for the UAG motif, and simultaneously an increased affinity for the CAG motif.

**Figure 6.**
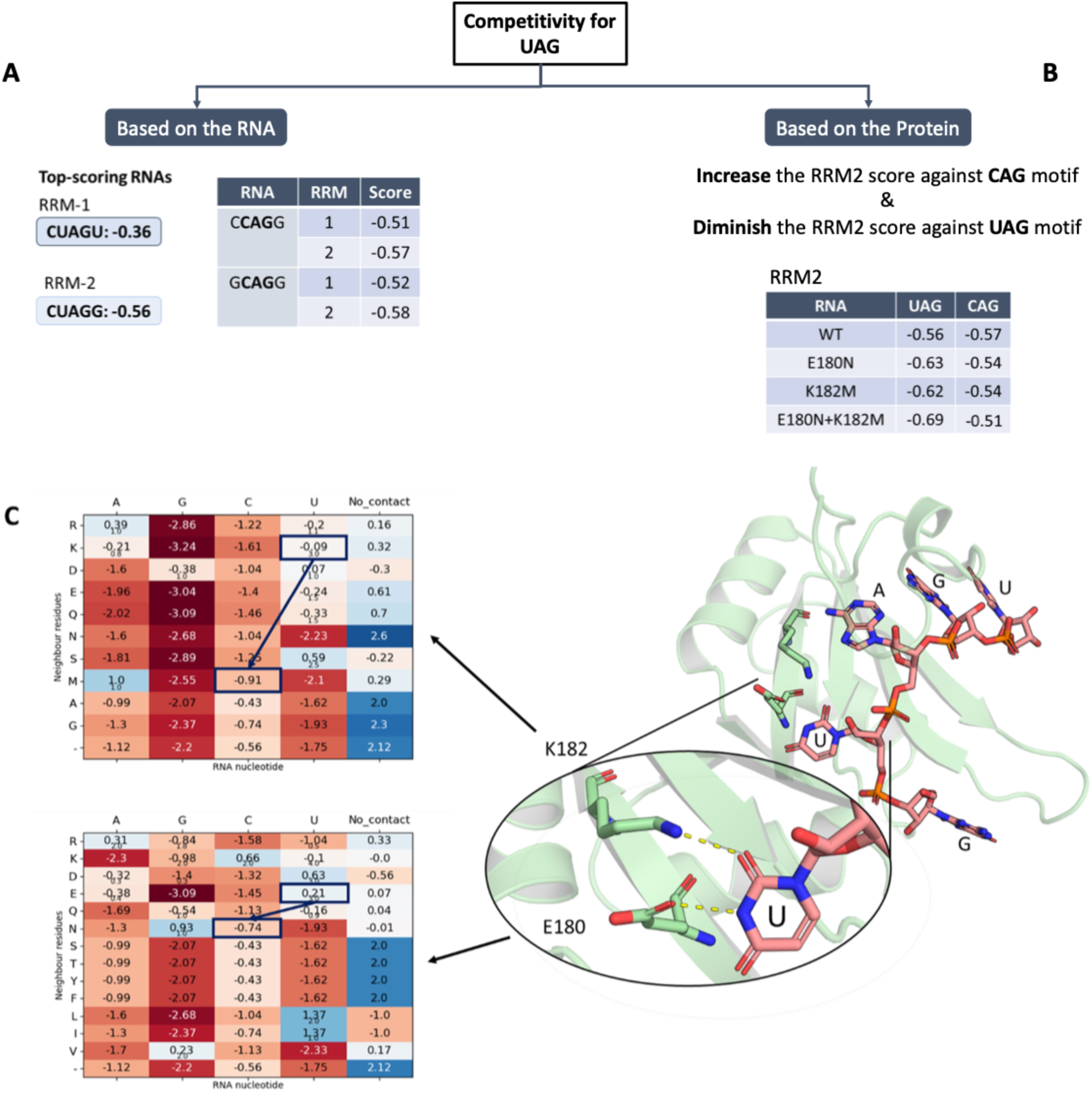
Scoring values from RRMScorer calculations. (A) Top-scoring RNAs for the isolated domains. (B) Top scores and mutations proposed based on the protein. (C) On the left, residues scoring values for the CAG motif in positions E180 and K182 of RRM-2. On the right, structural view of the second RNA Recognition Motif binding with GUAGU motif.

For this purpose, we analysed the possible residues involved in the recognition of the first pyrimidine of the 3-mer motif to switch the preference from uracil to cytosine (from UAG to CAG) (**Figure 6B**). The analysis of the x-ray structure of the RRM-2 in complex with GUAGU, already available on the Protein Data Bank (PDB ID: 5×3z), shows that E180 and K182 are involved in uracil recognition by engaging the amine and carbonyl groups of the nucleotide base, respectively (**Figure 6C**). According to the results provided by RRMScorer (**Figure 6B,C**) E180N and K182M are indeed the most promising mutations to switch the binding preference from an uracil towards a cytosine. The new amino acids present in the mutant maintain a similar overall steric bulk and are not expected to alter the RRM domain’s structure, while influencing the local charge distribution. Interestingly, these two residues are also present in the same position in other natural RRMs showing a preference for binding RNA oligonucleotides, bearing a cytosine in the same position as the chosen 3-mer motif.

### Experimental evaluation of protein mutant

First, the impact of each mutation on RRM-2 designed with RRMScorer was evaluated by analysing their binding to **oligo-L2** by SPR. The RRM-2 domain bearing the mutation E180N was called RRM2-M1 and the one containing the mutation K182M was called RRM2-M2. When binding **oligo-L2**, displaying an UAG motif, RRM2-M1 showed a slightly higher association rate constant and a two-fold faster dissociation rate constant than RRM-2, leading to a marginally lower affinity (**Figure 8, Table 2, Figures S7, S8**). In contrast, RRM2-M2 displayed an almost 7-fold slower association rate constant while keeping a similar dissociation rate constant, explaining the more than 5-fold weaker affinity.

After evaluating the effect of the single mutations on the binding, the impacts of the combined mutations for the MSI-1 RRM_1-2_ tandem domain on binding to the previously tested and designed RNA oligonucleotides were investigated. To evaluate if the mutations affect the affinity of the RRM-2 domain for the (G/A)U_1-3_AGU motif, the MSI-1 RRM_1-2_ tandem domain, bearing the two mutations E180N and K182M (MSI-1 RRM_1-2_-DM, hereafter), was titrated with increasing amounts of **oligo-L2** and NMR spectra were recorded. After the addition of **oligo-L2**, the signals of the free protein in the 2D ^1^H-^15^N TROSY spectrum decrease in intensity, while new cross-peaks, corresponding to the complex between MSI-1 RRM_1-2_-DM and **oligo-L2** appear and increase in intensity (**Figure 4**). More important, the signals of the free protein experiencing the largest decreases in intensity after the addition of RNA at a concentration of 25 µM to the protein solution (1:0.25 protein/RNA molar ratio), correspond all (but two) to residues located on RRM-1 domain (**Figure S11**). Although two residues in the C-terminal region of the RRM-2 domain still experience some effect, the two mutations on the RRM-2 domain largely shift the binding preference of **oligo-L2** towards the RRM-1 domain. However, the broadening of the signals suggests that some heterogeneity is still present at equimolar concentrations of protein and RNA. This is corroborated by SPR data where a biphasic mode of interaction is still present indicating that both RRM domains still bind **oligo-L2,** but the affinity of the bivalent interaction is more than 7-fold reduced when the double mutation is present (**Figure 8 and Table 3**). Also, InteractionMap analysis shows two peaks, representing the monovalent and bivalent interaction, but with a reduced affinity for the bivalent interaction of MSI-1 RRM_1-2_-DM compared to MSI-1 RRM_1-2_ (**Figure 7B**).

**Figure 7.**
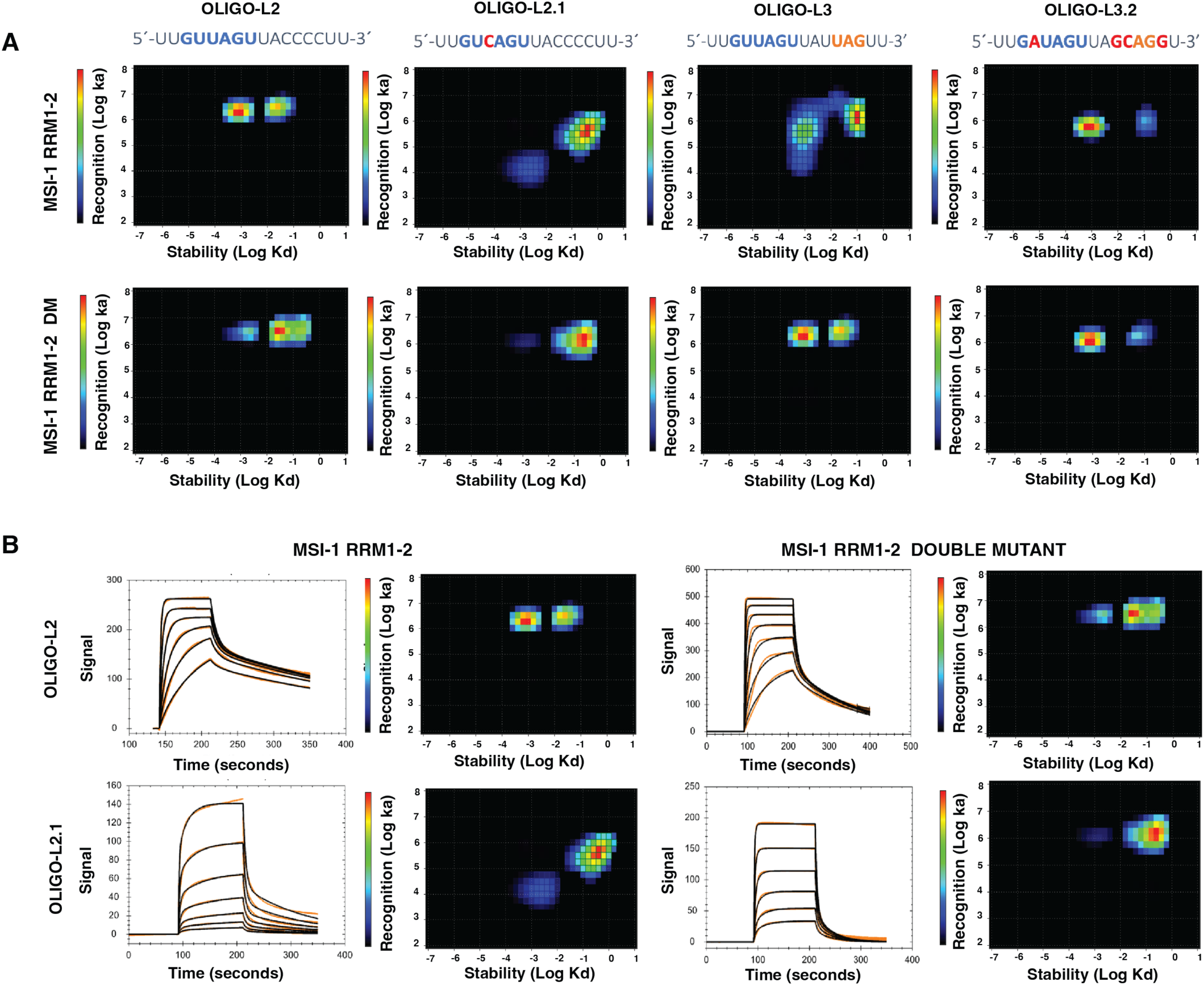
A) Representative InteractionMap from MSI-1 RRM_1-2_ and MSI-1 RRM_1-2_ DM interacting with **oligo-L2, oligo-L2.1, oligo-L3** and **oligo-L3.2**. B) Sensorgrams and InteractionsMaps, corresponding to interactions of MSI-1 RRM_1-2_ with **oligo-L2** with concentrations from 7.8 to 250 nM; MSI-1 RRM_1-2_ DM with **oligo-L2** with conc. ranging from 7.8 to 500nM; of MSI-1 RRM_1-2_ with **oligo-L2.1** with conc. ranging from 7.8 to 500 nM and MSI-1 RRM_1-2_ DM with **oligo-L2.1** with conc. ranging from 7.8 to 250nM.

**Figure 8.**
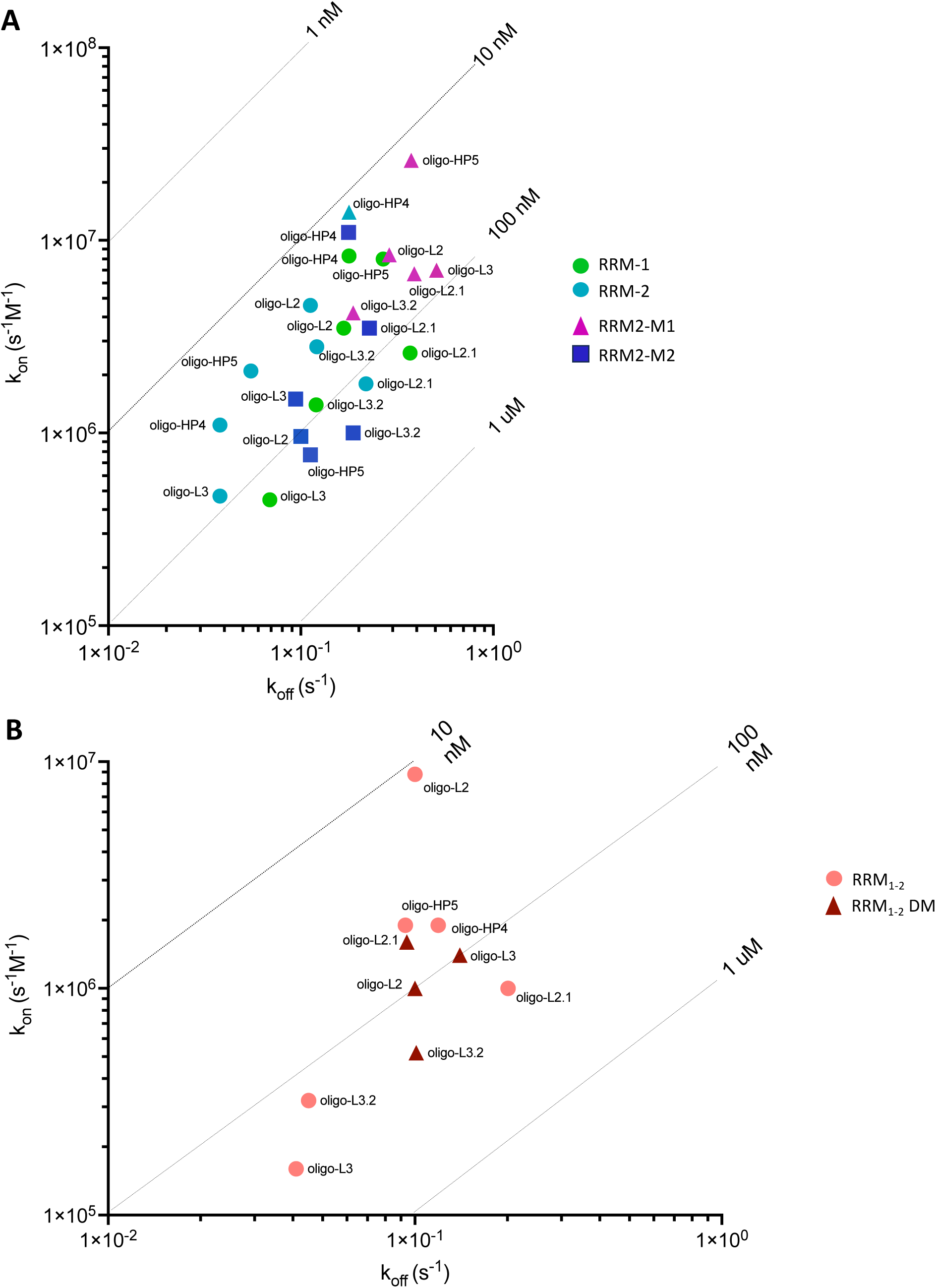
Isoaffinity kinetic plots of wild-type and mutants A) RRMs and B) tandem domains with the different oligos.

Then, the interaction of MSI-1 RRM_1-2_-DM with **oligo-L3** was analysed. Interestingly, after the addition of **oligo-L3**, at equimolar concentration with the protein, the linewidth of the new signals of the MSI-1 RRM_1-2_-DM in complex with **oligo-L3** is generally sharper than what was observed for the wild-type protein, suggesting the formation of a single species in solution for the protein/RNA complex (**see Suppl. Material**). More importantly, the signals mostly affected by interaction are assigned to residues majorly located on the RRM-1 domain, confirming the two mutations decrease the affinity of the RRM-2 domain towards the UAG motif (**see Suppl. Material**). When the interaction of MSI-1 RRM_1-2_-DM with **oligo-L3** was characterized with SPR, the monovalent interaction had a 2.5 stronger affinity value, due to a 10-fold faster association rate compared with the non-mutated tandem binding (**Figure 8 and Table 3**). The monovalent interaction of MSI-1 RRM_1-2_-DM with **oligo-L3** is also very similar to that with **oligo-L2,** indicating that the binding is no longer affected by the competition of the two RRM domains for the same motif, as observed for the interaction of the wild-typed protein with **oligo-L3**. This was confirmed by InteractionMap analysis, which showed less heterogenicity with smaller and better-defined peaks when compared with the wild-type protein (**Figure 7A**). Nevertheless, MSI-1 RRM_1-2_-DM still has a relatively high affinity for the bivalent interaction which is less than 2-fold weaker compared to the interaction of the wild-type with **oligo-L2,** but more than 4-fold stronger than the MSI-1 RRM_1-2_-DM interaction with **oligo-2** (**Figure 7, 8 and Table 3**).

Multiple interactions were also observed by NMR when MSI-1 RRM_1-2_ DM was titrated with **oligo-HP4**. In this case, more than one species seems to be still present in solution, as suggested by the broadening of the signals (**Figure 4 and Suppl. Figure S14**). Nevertheless, in the presence of **oligo-HP4**, a lower heterogeneity is observed for the double mutant compared to the wild-type protein, as described in the **Supplementary Material**. However, as for wild-type MSI-1 RRM_1-2_ the opening of the hairpin structure may also be responsible for the observed conformational heterogeneity.

### Experimental evaluation of the RNA mutants

To evaluate the preference of binding for the minimal RNA motif, we designed **oligo-L2.1** (5’-UU**GUCAGU**UACCCCUU −3’), where the UAG motif has been replaced by a CAG motif. When interacting with the isolated RRM-1 and RRM-2 domains, SPR data revealed a 3-fold and almost 5-fold weaker affinity, respectively. Conversely, RRM2-M1 and RRM2-M2, bearing mutations to shift the binding preference from UAG to CAG, showed a slightly (3-fold and less than 2-fold) stronger affinity with respect to wild-type RRM-2 (see **Figure 8, Table 2 and Figure S8**), mostly due to a better recognition. These effects are more evident when the **oligo-L2.1** interaction with MSI-1 RRM_1-2_ tandem domain is evaluated. **Oligo-L2.1,** is expected to diminish the binding partitioning between the two domains and to exhibit a lower binding affinity than **oligo-L2** for MSI-1 RRM_1-2_. This is confirmed by the 4-fold weaker affinity for monovalent binding. The bivalent interaction, although still present, provides a small contribution to the overall interaction. Therefore, only the values for the monovalent interaction were shown in **Table 3**. This negligible contribution was confirmed by InteractionMap analysis, which did not show a clear stabilised fraction compared to the previously obtained results with **oligo-L2** (**Figure 7A**). This indicates that the MSI-1 RRM_1-2_ reduces its ability to form bivalent interactions on the sensor chip for the CAG binding motif.

Compared to binding of MSI-1 RRM_1-2_ to **oligo-L2.1**, the mutated tandem protein showed a more than 3-fold increase in affinity for the monovalent interaction with improved recognition and slower dissociation. The affinity of MSI-1 RRM_1-2_ DM for **oligo-L2.1** is in the same range of that of the wild-type protein for **oligo-L2**, **-HP4** and **-HP5** (**Figure 8 and Table 3**). As previously observed for the wild-type protein, the interaction is strongly dominated by a rapid 1:1 interaction and became similar to that observed for interactions of isolated RRMs with the oligo. The contribution of bivalent interactions to the overall sensorgrams is low at higher oligo immobilization levels becomes neglectable at lower oligo densities. Therefore, binding curves were analysed using the 1:1 model for two replicates with lower RNA densities while the 1:2 model was applied to interactions at higher RNA densities and only the results from the strongly dominating rapid interaction was reported in **Table 3**. The weak or insignificant contribution of bivalent interaction indicates that RRM-2 is leading the recognition while RRM-1 interaction with **oligo-L2.1** is weak. This was confirmed by InteractionMap analysis of the MSI-1 RRM_1-2_ DM interaction with **oligo-L2.1** that shows a dominating peak corresponding to the monovalent interaction (**Figure 7A**).

NMR spectroscopy was used to investigate the interaction of the modified RNA oligonucleotides bearing two binding sites (**oligo-L3.2** (5’-UU**GAUAGU**UAG**CAG**GU-3’) and **oligo-HP4.2** (5’-AAGC**GAUAGU**UAUG**CAG**GCGCUU-3’)) with the wild-type MSI-1 RRM_1-2_ protein and with the double mutant MSI-1 RRM_1-2_-DM. The interaction of the wild-type tandem domain protein with **oligo-L3.2** is in the slow exchange regime on the NMR timescale, as observed with the original **oligo-L3** (see **Suppl. Material**). However, at equimolar protein/RNA ratio, the signals of the wild-type MSI-1 RRM_1-2_ in the 2D ^1^H-^15^N TROSY spectrum appear split, suggesting the presence of multiple species in solution (**Figure 4**). This effect is more evident than with the original **oligo-L3**, and it is not possible to rule out that one of the two RRM domains is not interacting with the RNA. As expected, the modification of the RNA binding site weakens the interaction of the oligo with the wild-type MSI-1 RRM_1-2_ protein (see **Supplementary Material**). In this case, residues experiencing the largest effect are spread on both domains and not only on RRM-1, as observed for the interaction of the double mutant with **oligo-L3**. SPR experiments showed a 2-fold faster association of MSI-1 RRM_1-2_ the two isolated RRM domains to **oligo-L3.2** compared with **oligo-L3**, resulting in a two-fold stronger affinity indicating that the CAG nutation reduced heterogeneous recognition induced by binding to multiple motifs on the same RNA strand. Conversely, the double mutant exhibits similar affinities for the monovalent and bivalent interaction with **oligo-L3** and **oligo-L3.2.** As expected, the double mutant MSI-1 RRM_1-2_ DM binds 2.6-fold slower to **oligo-L3.2** compared to **oligo-L3**. However, the association is still more than three times faster than that observed for the interaction between the wild-type tandem domain and **oligo-L3** (**Table 3**) and InteractionMap analysis for both the wild-type and mutated tandem domain with **oligo-L3.2** showed well-defined peaks similar to those observed for **oligo-L2** (**Figure 7A**). This is supported by NMR data that show that, in the presence of **oligo-L3.2** in a 1:1 molar ratio, a single species is visible for MSI-1 RRM_1-2_-DM (**Figure 4**).

The effects of **oligo-HP4.2** on the wild-type MSI-1 RRM_1-2_ and MSI-1 RRM_1-2_-DM were also investigated with NMR spectroscopy (see **Supplementary Material**). Modifications on the RNA construct have weakened the interaction of the RRMs for the second binding site in order to disfavour the formation of possible species involving more than a single MSI-1 protein. Unfortunately, heterogeneity is still observed in solution with different combinations of protein-RNA binding modes: e.g. complexes with RRM-1 or RRM-2 bound to the G/AU_1-3_AGU binding site for both the wild-type and double mutated proteins.

These findings corroborate the idea that the modifications present in **oligo-HP4.2**, when combined with mutations on the protein, do not prevent the formation of different complexes in which the (G/A)U_1-3_AGU motif can interact with either one or the other RRM domain.

## Discussion

The design of novel RRMs or RNA strands with high affinity could be relevant to investigating the regulation of gene expression, as well as in the discovery of *in vivo* RNA targets of RRMs with a still unknown function or interaction. In this study, we have used the computational tool RRMScorer to design a set of mutants of the human Musashi-1 protein able to bind a novel RNA sequence to modify the binding selectivity. Using complementary biophysical techniques, we have investigated the contributions to the interaction between the wild-type and protein MSI-1 mutants with the known and redesigned RNA strands to gain insight on how they interact, and which are the key structural features involved in the interaction.

According to our results, the two isolated domains of human Musashi-1 behave similarly when interacting with oligonucleotides like **oligo-L2,** bearing a single motif (G/AU_1-3_AGU), for which both domains show nanomolar affinity. This finding is supported by the similar kinetic profile observed for the isolated domains and by NMR and kinetics experiments performed on the tandem domain in the presence of **oligo-L2.** Indeed, RRM-1 and RRM-2 compete for this site, and a biphasic dissociation is observed for the tandem domain. This kinetic profile is related to the ability to rapidly and stably bind to two different oligos at high oligo/MSI-1 ratios. Interesting are also the results obtained with **oligo-L3** which contains two recognition sites. The capability of this oligo to bind two RRM-1 domains to form a 2:1 RRM-1/RNA complex is proved by the NMR data and the SEC-MALS analysis, as well by SPR data. The slower association rate constant (k_a1_) for both RRM-1 and RRM-2 when binding **oligo-L3**, compared with the other oligos, could be due to steric hindrance, with the binding of a second RRM hampered by the already bound RRM. For both **oligo-HP4** and **-HP5,** also harbouring two UAG binding motifs, the low amount of two different RRMs bound to the same RNA strand, probably due to the hairpin conformation, seems to be responsible for the fast association. This finding is in agreement with the formation of 1:1 protein/**oligo-L3** complexes in the presence of the tandem domain. The ability of both domains being able to recognize the RNA is also reflected in the similar association rates obtained from the isolated RRMs interacting with **oligo-L2**.

This study has also provided important information on the influence of the RNA structure on the recognition process and binding by MSI-1. The NMR and SPR data show that the two domains bind the hairpin with high affinity, as also observed for the linear strands. More importantly, the SEC-MALS analysis shows that two isolated RRM-1 can bind **oligo-HP4** at the same time, thus providing a 2:1 protein/RNA complex. Overall, the data collected on **oligo-HP4** show that: i) the presence of the binding sites in the loop of numb mRNA does not inhibit binding; ii) the position of the two binding sites within this mRNA strand allows it to accommodate two RRM domains at the same time and iii) binding kinetics of both RRMs are similar and therefore nor the presence of the loop nor the position of the binding site affect the kinetic profiles. Further interesting information was obtained from the analysis of the experimental data collected on the tandem domain in the presence of **oligo-HP4**. The general decrease in signal intensity observed for the cross-peaks of the free protein (stronger for the residues of the protein binding sites) during the NMR titration, without the appearance of new signals, suggests competition between the two domains for the two binding sites present in the hairpin. However, in contrast to what was previously observed with **oligo-L2**, the general decrease in signal intensity indicates the presence of high molecular weight species involving two or more MSI-1 proteins. Partial unfolding of the hairpin upon MSI-1 binding is also a possible explanation for the heterogeneity of the conformations sampled in solution by the protein/RNA complexes, as suggested by fluorescence quenching assays and ^1^H imino NMR spectra. More importantly, these findings seem to rule out the capability of the two RRMs belonging to a single MSI-1 to bind the two sites of a hairpin structure at the same time, as observed instead with the linear RNA, **oligo-L3**. The formation of these intermolecular complexes over intramolecular complexes is backed by the SPR results as previously described ^30^. The influence of RNA coating density on the stabilised bivalent fraction suggested that intermolecular binding is preferred. This intermolecular binding of MSI-1 RRM_1-2_ to different RNA strands is also suggested by the well-defined peaks obtained with InteractionMap analysis (**Figure 7**).

Interesting is also the behaviour of MSI-1 in the presence of the **oligo-HP5**. The high affinity of the isolated domains for this hairpin is proven by the NMR and SPR data. Also, the tandem domain binds the **oligo-HP5** with high affinity, but the evolution of the NMR spectra upon the addition of the hairpin suggests a further different behaviour. The persistence of the signals of the free protein together with the signals of the new species, even in the presence of an excess of the hairpin, raises several questions about how MSI-1 interacts with this oligo. One possibility could be that the hairpin is only bound at one site (perhaps the G/AU_1-3_AGU site on the loop) of the oligo, and that its interaction with the second domain is prevented by the involvement of the second site (UAG) in the double strand structure. Furthermore, following this hypothesis, the structure of the MSI-1-RNA complex could make the second domain of the protein unavailable to bind a second oligo. Unfolding of the hairpin structure upon binding of the tandem domain is also observed for **oligo-HP5**.

The data collected on the wild-type human MSI-1 tandem domain thus show that the protein can bind the consensus binding motif G/AU_1-3_AGU as well as the UAG sequence when present in linear RNA sequences or folded RNA structures. The protein binds all four investigated oligos with high affinity (**Figure 8**, **Table 2 and 3**). However, based on the NMR data, it appears that only the **oligo-L3** seems to be able to interact with both the RRM-1 and RRM-2 of a single MSI-1 simultaneously, forming 1:1 protein-RNA complexes. This observation is further supported by the kinetic profile of the isolated RRMs binding to **oligo-L3**, particularly due to the slower association rate constant. The InteractionMap analysis revealed broad peaks, suggesting a heterogeneous recognition, which can be attributed to steric hindrance between two domains interacting simultaneously on the same strand. This hindrance is likely due to a fast binding of the first RRM and the slower binding of the second RRM domain. This interpretation is supported by the data obtained from investigating the interaction of the tandem protein with **oligo-HP4** and **-HP5,** both of which have two binding motifs displaying clear peaks in the InteractionMap analysis (**Figure 7**).

The presence of multiple RNA binding domains in RNA-binding proteins (RBPs) allows them to recognize specific RNA sequences with high affinity. This is achieved through the engagement of different consensus binding motifs in the same strand, and the cooperation of the RNA recognition motifs (RRMs). ^3–5,34–39^. In other cases, RRMs can bind RNA independently, increasing the chance for the protein to encounter its specific RNA binding sequence. Interestingly, our study of the tandem domain of human Musashi-1 protein has revealed a complex and dynamic mode of interaction, where the isolated domains of a tandem RRM protein compete for the same binding site, instead of cooperating to stabilize the interaction. This highlighted the relevance of Musashi-1 not only as a potential pharmacological target but also as a model for investigating the molecular basis of RNA recognition by RBPs. Additionally, it validates the RRMScorer predictor for the design of new RNA and protein mutants.

In this regard, the results obtained from the protein mutant and modified oligonucleotides designed by RRMScorer are equally significant and intriguing. In the presence of **oligo-L2**, MSI-1 RRM_1-2_ DM exhibits in the monovalent and bivalent interaction almost a 2-fold and a 10-fold lower affinity, respectively, for the (G/A)U_1-3_AGU motif. MSI-1 RRM_1-2_ DM showed a stronger monovalent interaction to **oligo-L2.1**, compared with the binding of both MSI-1 RRM_1-2_ and MSI-1 RRM_1-2_ DM to **oligo-L2**. This is mostly due to the improved affinity of mutated RRM-2 for the CAG motif, as expected from the RRMScorer prediction. The bivalent interaction, already weak in MSI-1-RRM_1-2_ DM with **oligo-L2**, is completely absent with **oligo-L2.1**.

NMR data acquired on MSI-1 RRM_1-2_ DM with **oligo-L3** further corroborate the effect of the mutations in the reduction of competition of RRM-2 with RRM-1, for the UAG motif. Although no major differences in affinity are observed, the NMR titration of MSI-1 RRM_1-2_ DM with **oligo-L3** shows the presence of a single MSI-1/RNA complex with RRM-1 mostly affected by spectral changes.

Conversely, with the wild-type MSI-1 the competition between RRM-1 and RRM-2 for the two UAG sites leads to the formation of two or more species. A better selectivity for RRM-1 was also observed when the MSI-1 RRM_1-2_ DM was titrated with the folded **oligo-HP4**. The decreased affinity of RRM-2 for the UAG motif leads to weak effects on the RRM-2 domain and to a reduction of heterogeneity in solution, even though multiple species are still observed.

It is important to highlight that, the CAG mutation on the two RNA strands restores the affinity of the mutated RRM-2 domain for the two oligos leading to a more specific binding compared to when the wild-type MSI-1 was titrated with non-mutated RNA strands by NMR. In particular, a single species is obtained when a stoichiometric amount of **oligo-L3.2** was added to the MSI-1 RRM_1-2_ DM protein. SPR results showed that the mutated protein had a better recognition for **oligo-L3.2**, resulting in slightly higher affinity than the wild-type tandem protein. This confirms the results from NMR and, together with the results obtained for MSI-1 RRM_1-2_ DM interacting with **oligo-L2.1**, showed that MSI-1 RRM_1-2_ DM has improved recognition for the CAG motif, as observed from a faster on rate, compared to the non-mutated protein. The improved selectivity obtained with protein mutations and RNA modification observed in NMR is also supported by the results obtained with the hairpin RNA constructs. Two predominant species seem to be present in solution when **oligo-HP4.2** is added to MSI-1 RRM_1-2_ DM protein. This behaviour is significantly different from that of the wild-type MSI-1 in the presence of non-mutated **oligo-HP4**, where a larger heterogeneity was observed.

Finally, SPR experiments allowed us to investigate the individual contribution of each of the mutations in the RRM-2 domain to the binding kinetics and affinity. RRM2-M1, with E180N mutation, showed a higher affinity than RRM2-M2. When analysing the RRM-2 interaction to **oligo-L2.1**, that contains no UAG motif and a single CAG motif, data show that RRM-2 mutations increase the affinity that is mainly affected by the E180N mutation.

Summarizing, this study provides a comprehensive analysis of human Musashi-1 and offers valuable insights into the recognition and binding of RNA by RBPs, suggesting this protein as a promising candidate to develop mutants with high selectivity for specific RNA sequences for in vitro and in vivo studies and applications. The findings validate the reliability of the approach designed to enhance the specificity of protein-RNA interactions, and the quality of the prediction provided by RRMScorer, paving the way for the development of new tools for biological investigations, and potential biotherapeutics.

## Acknowledgements

This work has been supported by the Marie Skłodowska-Curie Innovative Training Network (MSCA-ITN) RNAct supported by European Union’s Horizon 2020 research and innovation programme under grant agreement No 813239, the Regione Toscana (CERM-TT, BioEnable and PANCREAS-AD bando salute 2018), the Italian Ministero dell’Istruzione, dell’Università e della Ricerca through PRIN 2017A2KEPL, the “Progetto Dipartimenti di Eccellenza 2018-2022” to the Department of Chemistry “Ugo Schiff” of the University of Florence, the Recombinant Proteins JOYNLAB laboratory, and the project FISR2021_SYLCOV. The authors acknowledge the support and the use of resources of Instruct-ERIC, a landmark ESFRI project, and specifically the CERM/CIRMMP Italy centre. We acknowledge also the project “Potentiating the Italian Capacity for Structural Biology Services in Instruct Eric (ITACA.SB)” (Project n° IR0000009) within the call MUR 3264/2021 PNRR M4/C2/L3.1.1, funded by the European Union NextGenerationEU. This study was also supported by the Next Generation EU project within the MUR PNRR “National Center for Gene Therapy and Drugs based on RNA Technology” (Project No. CN00000041 CN3 RNA).

## Author contributions

M.F., T.M, and A.P.R. conceptualized the study. A.P.R cloned expression plasmids, expressed and purified proteins. L.C and A.P.R planned, executed, and analysed NMR experiments. A.P.R. carried out SEC-MALS analysis. L.S planned and conducted fluorescence quenching assays and NMR spectroscopy for quenching assays under the supervision of M.S.. G.P.R designed, performed, and analysed kinetic experiments with support of R.A.H, under the supervision of J.B., U.H.D and W.H.. J.R.M and A.P.R planned and J.R.M executed and analysed RRMScorer predictions under the supervision of W.V.. Together, M.F., A.P.R, L.C., G.P.R. and T.M, wrote the manuscript with input from other authors.

